# Serotonin neurons modulate learning rate through uncertainty

**DOI:** 10.1101/2020.10.24.353508

**Authors:** Cooper D. Grossman, Bilal A. Bari, Jeremiah Y. Cohen

## Abstract

Regulating how fast to learn is critical for flexible behavior. Learning about the consequences of actions should be slow in stable environments, but accelerate when that environment changes. Recognizing stability and detecting change is difficult in environments with noisy relationships between actions and outcomes. Under these conditions, theories propose that uncertainty can be used to modulate learning rates (“meta-learning”). We show that mice behaving in a dynamic foraging task exhibit choice behavior that varied as a function of two forms of uncertainty estimated from a meta-learning model. The activity of dorsal raphe serotonin neurons tracked both types of uncertainty in the foraging task, as well as in a dynamic Pavlovian task. Reversible inhibition of serotonin neurons in the foraging task reproduced changes in learning predicted by a simulated lesion of meta-learning in the model. We thus provide a quantitative link between serotonin neuron activity, learning, and decision making.

---

Models from control theory and reinforcement learning (RL) propose that behavioral policies are learned through interactions between the nervous system and the environment (Bertsekas and Tsitsiklis, 1996; Sutton and Barto, 1998). Within this framework, an animal learns from discrepancies between expected and received outcomes of actions (reward prediction errors, RPEs). The rate at which learning occurs is usually treated as a fixed parameter, but optimal learning rates vary when the environment changes. Consequently, animals should vary how rapidly they learn in order to behave adaptively and maximize reward. Normatively, learning rates should vary as a function of uncertainty (Dayan et al., 2000; Yu and Dayan, 2005; Soltani and Izquierdo, 2019). When some amount of uncertainty is expected (also referred to as outcome variance or risk), learning rates should decrease (Preuschoff et al., 2006; Preuschoff and Bossaerts, 2007; Diederen and Schultz, 2015). Slower learning helps maximize reward when relationships between actions and outcomes are probabilistic but stable. This prevents animals from abandoning an optimal choice due to short-term fluctuations in outcomes. However, it is also important to detect changes in the underlying statistics of an environment. Here, deviations from expected uncertainty (“unexpected uncertainty”) should increase learning rates (Yu and Dayan, 2005; Payzan-LeNestour and Bossaerts, 2011; Payzan-LeNestour et al., 2013; O’Reilly, 2013; Faraji et al., 2018). Tuning decision making in this way is known as “meta-learning,” and there is evidence that humans and other animals use this strategy (Behrens et al., 2007; Herzfeld et al., 2014; Massi et al., 2018; Soltani and Izquierdo, 2019). How does the nervous system control how rapidly to learn from recent experience?

Several theories propose that neuromodulatory systems enable meta-learning (Daw et al., 2002; Doya, 2002; Yu and Dayan, 2005). One such system comprises a small number of serotonin-releasing neurons (on the order of 10^4^ in mice; Ishimura et al., 1988) with extensive axonal projections. This small group of cells affects large numbers of neurons in distributed regions (Steinbusch, 1981; Jacobs and Azmitia, 1992; Ren et al., 2018; Awasthi et al., 2020) that are responsible for learning and decision making. The activity of these neurons changes on behaviorally-relevant timescales—both fast (hundreds of milliseconds) and slow (tens of seconds; Liu et al., 2014; Cohen et al., 2015; Li et al., 2016; Matias et al., 2017). Serotonin receptor activation can induce short-term changes in excitability (Andrade, 2011; Celada et al., 2013) as well as long-lasting synaptic plasticity (Lesch and Waider, 2012).

Prior research demonstrates that serotonin neurons modulate flexible behavior in changing environments (Clarke et al., 2004, 2007; Boulougouris and Robbins, 2010; Bari et al., 2010; Brigman et al., 2010; Matias et al., 2017; Iigaya et al., 2018). Serotonin axon lesions (Clarke et al., 2004, 2007) or reversible inactivation of dorsal raphe serotonin neurons (Matias et al., 2017) impaired behavioral adaptation to changes in action- or stimulus-outcome mappings. Importantly, in these experiments, animals were still capable of adapting their behavior, but did so more slowly. Conversely, brief excitations of serotonin neurons in a probabilistic choice task enhanced learning rates after long intervals between outcomes (Iigaya et al., 2018). These studies show that serotonin neurons modulate how quickly an animal adapts to a change in causal relationships in the environment. Thus, serotonin neurons may guide learning using the statistics of recent outcomes. However, a mechanistic understanding of the relationship between serotonin neuron activity and meta-learning has not been established.

We designed a dynamic foraging task for mice and recorded action potentials from dorsal raphe serotonin neurons. We developed a generative model of behavior by modifying an RL model to include meta-learning. Adding meta-learning to the model captured a unique feature of observed behavior that a model of behavior with a fixed learning rate could not explain. We found that the activity of approximately half of serotonin neurons correlated with the “expected uncertainty” variable from the model on long timescales (tens of seconds to minutes) and “unexpected uncertainty” at the time of outcome. Simulated removal of meta-learning from the model predicted specific changes in learning that were reproduced by chemogenetic inhibition of dorsal raphe serotonin neurons. Thus, we demonstrate a quantitative link between serotonin neuron activity and behavior.

## Mice display meta-learning during dynamic decision making

We trained thirsty, head-restrained mice (21 female, 27 male) on a dynamic foraging task in which they made choices between two alternative sources of water (Bari et al., 2019). Sessions consisted of about 300 trials (280 ± 66.6) with forced inter-trial intervals (1–31 s, exponentially distributed). Each trial began with an odor “go” cue that informed the animal that it could make a choice, but otherwise gave no information (Figure 1a,b). During a response window (1.5 s) the mouse could make a decision by licking either the left or the right spout. As a consequence of their choice, water was delivered probabilistically from the chosen spout. The reward probabilities (*P* (*R*)) assigned to each spout changed independently and randomly, in blocks of 20–35 trials (drawn from a uniform distribution). These reward contingencies were drawn from a set of three probabilities (either *P* (*R*) ∈ {0.1, 0.5, 0.9} or *P* (*R*) ∈ {0.1, 0.4, 0.7} for a given mouse) and were not signaled to the animal.

**Figure 1.**
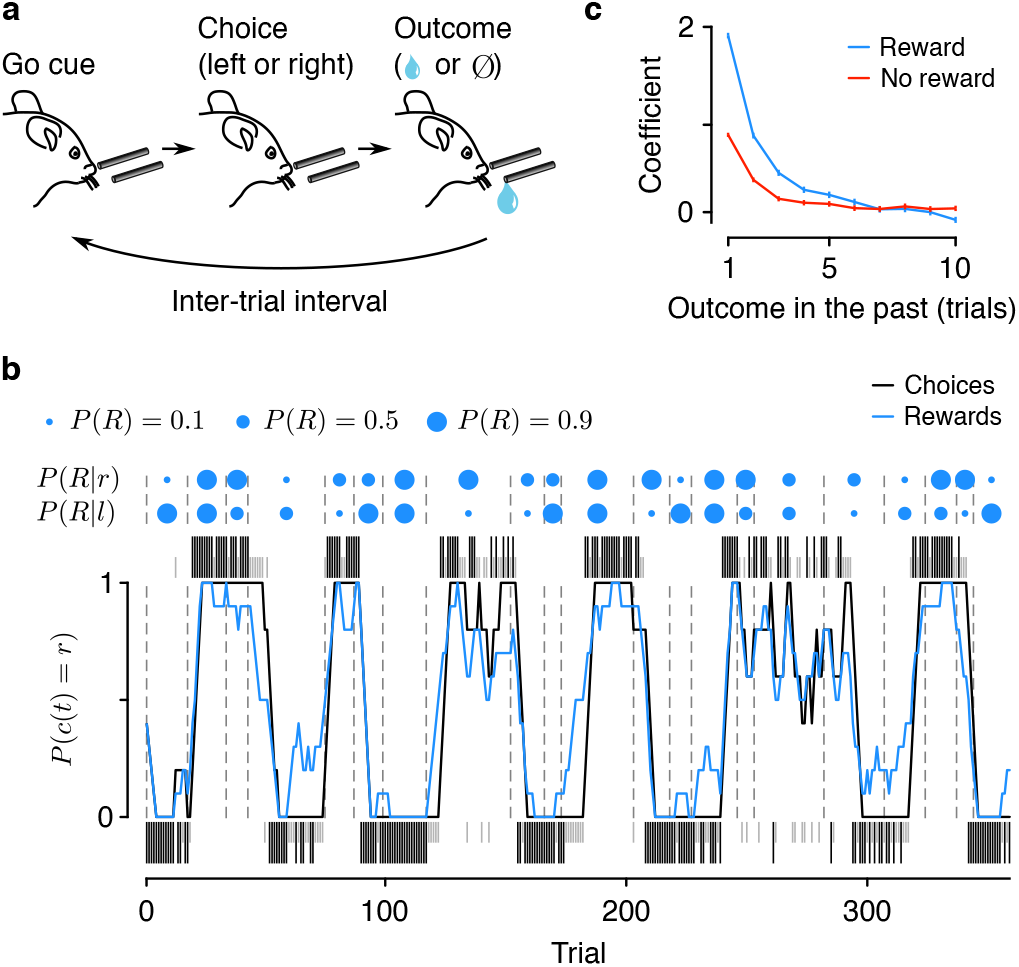
Mice forage dynamically for rewards. (a) Dynamic foraging task in which mice chose freely between a leftward and rightward lick, followed by a reward with a probability that varied over time. (b) Example mouse behavior from a single session in the task. Black (rewarded) and gray (unrewarded) ticks correspond to left (below) and right (above) choices. Black curve: mouse (smoothed over 5 trials, boxcar filter) choices. Blue curve: rewards (smoothed over 5 trials, boxcar filter). Blue dots indicate left/right reward probabilities and dashed lines indicate a change in reward probability (*P* (*R*)) for at least one spout. (c) Logistic regression coefficients for choice as a function of outcome history. Error bars: 95% CI.

Mice mostly chose the higher-probability spout (Figure A1a; correct rate, 0.68 ± 0.029) and harvested rewards (reward rate, 0.57 ± 0.021 rewards trial^−1^) over many sessions (13.8 ± 6.93 sessions mouse^−1^). We first fit statistical models to quantify the effect of outcome history on choice. These logistic regressions revealed that mice used experience of recent outcomes to drive behavior (Figure 1c; time constants, 1.31±0.25 trials for rewards, 1.04±0.13 trials for no rewards, 95% C.I.). Similarly, we quantified the effect of outcomes on the latency to make a choice following the go cue. Consistent with previous findings (Bari et al., 2019), this model demonstrated a large effect of recent rewards on speeding up response times (Figure A1b,c; time constant, 1.76 ± 0.27 trials, 95% C.I.).

These statistical findings indicate that mice dynamically learned from recent experience. To understand the nature of this learning, we constructed a generative model from a family of RL models called *Q*-learning (Bertsekas and Tsitsiklis, 1996; Sutton and Barto, 1998). This class of models creates a behavioral policy by maintaining an estimate of the value of each action (the expected reward from making that action). Using these values to make choices, the model then learns from those choices by using the RPE to update the action values, thereby forming a new policy (Figure 2a). How much to learn from RPEs is determined by the learning rate parameters. While these parameters are typically fixed across behavior, they need not be constant; they could vary according to statistics of the environment (meta-learning; Behrens et al., 2007).

**Figure 2.**
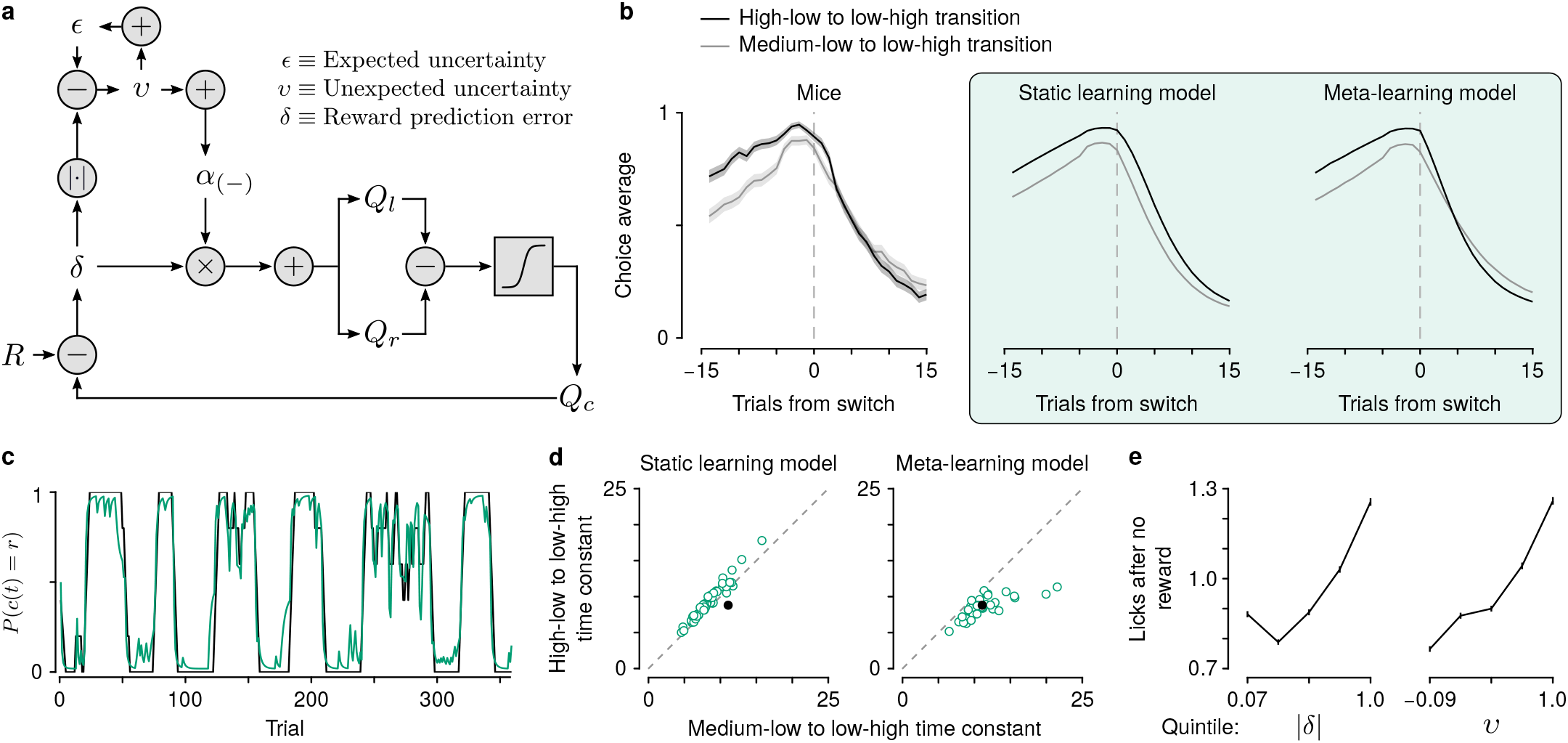
Mice learn at variable rates. (a) Schematic of the meta-learning model algorithm. *Relative value* (*Q*_*r*_ − *Q*_*l*_) is used to make choices. The predicted value of a choice (*Q*_*c*_) is compared to reward (*R*), to generate a *reward prediction error* (*δ*). *|δ|* is used to update *expected uncertainty* (*ϵ*), which is compared with the prediction error to generate *unexpected uncertainty* (*v*). *v* then determines how rapidly to learn from *δ*, thereby updating *Q*_*r*_ and *Q*_*l*_. (b) Left: actual mouse behavior at transitions in which reward probabilities change simultaneously (*n* = 225 high-low to low-high, *n* = 236 medium-low to low-high). Lines are mean choice probability relative to the spout that initially has the higher probability, shading is Bernoulli SEM. Middle: simulated behavior at transitions using static learning model parameters fit to actual behavior. Right: simulated behavior at transitions using meta-learning model parameters fit to actual behavior. (c) Estimated choice probability of actual behavior (black, same as Figure 1b) and choice probability estimated with the meta-learning model (green) smoothed over 5 trials (boxcar filter). (d) Time constants from exponential curves fit to simulated choice probabilities (like those shown in (b)) for each mouse (*n* = 48, green circles) compared to the actual mouse behavior (black circle). Left: static-learning model (probability that mouse data come from simulated data distribution, *p <* 10^−6^). Right: meta-learning model (*p* = 0.89). (e) Spout licks following no reward as a function of *δ* from the static learning model (left, regression coefficient = 0.38, *p <* 10^−20^) or *ϵ* from the meta-learning model (right, regression coefficient = 0.45, *p <* 10^−20^).

We first fit a model to mouse behavior in which learning rates were constant. The model included separate parameters for learning from positive and negative RPEs because learning from rewards and no rewards was demonstrably asymmetric (Figure 1c), consistent with previous reports (Lefebvre et al., 2017; Dorfman et al., 2019; Dabney et al., 2020). This model fit overall behavior well (Bari et al., 2019), but was unable to capture a specific feature of behavior around transitions in reward probabilities (Figure 2b). In rare instances, both reward probabilities were reassigned within 5 trials of each other. When the probability assignments flipped from high and low to low and high (for example, from 0.9 on the left and 0.1 on the right to 0.1 on the left and 0.9 on the right), mice rapidly shifted their choices to the new higher-probability alternative. However, when reward probabilities transitioned from medium and low to low and high (for example, from 0.5 on the left and 0.1 on the right to 0.1 on the left and 0.9 on the right), mice took longer to adapt to the change (Figure 2b; effect of trial from transition *F*_1,28_ = 217, *p <* 10^−13^ and trial from transition × transition type interaction *F*_1,28_ = 5.23, *p* = 0.030, linear mixed effects model). This difference in choice adaptation was still apparent when choice probabilities prior to the transition were identical (Figure A2a), demonstrating that the difference in outcome history is responsible for this effect on choice adaptation.

Based on outcome history, the transition from high to low is more obvious than the transition from medium to low. This observation is consistent with learning rates varying as a function of how much outcomes deviate from a learned amount of variability (expected uncertainty). Thus, we designed a model (Figure 2a) that learns an estimate of the expected uncertainty of the behavioral policy by calculating a moving, weighted average of unsigned RPEs (Soltani and Izquierdo, 2019). Increases in expected uncertainty cause slower learning. This computation helps maximize reward when outcomes are probabilistic but stable (Dayan et al., 2000; Preuschoff and Bossaerts, 2007; Diederen and Schultz, 2015). The model then calculates the difference between expected uncertainty and unsigned RPEs (unexpected uncertainty), integrating over trials, to determine how quickly the brain learns from those outcomes (Krugel et al., 2009; Payzan-LeNestour and Bossaerts, 2011; Payzan-LeNestour et al., 2013; Faraji et al., 2018). Intuitively, large RPEs that differ from recent history carry more information because they may signal a change in the environment and should therefore enhance learning.

When we modeled the mouse behavior with meta-learning in this way, simulations using fitted parameters reproduced the transition behavior (Figure 2b-d). It was only necessary to modulate learning from negative RPEs to capture the behavior of mice around these transitions, perhaps due to the asymmetric effect of rewards and no rewards on behavior (Figure 1c). Interestingly, not all forms of meta-learning were capable of mimicking mouse behavior. We were unable to reproduce the observed behavior using a model previously proposed to modulate learning rates and explain serotonin neuron function (Figure A2c,d; Daw et al., 2002; Wittmann et al., 2020). A Pearce-Hall model (Pearce and Hall, 1980), which modulates learning as a function of RPE magnitude in a different way, was also unsuccessful (Figure A2c,d).

To capture this transition behavior, our meta-learning model leveraged a higher learning rate following high-low to low-high transitions than following medium-low to low-high. Prior to the transitions, expected uncertainty was lower when the animal was sampling the high probability spout as opposed to the medium probability spout (Figure A2e, *t*_473_ = 11.8, *p <* 10^−27^, paired *t*-test). When the reward probabilities changed, the deviation from expected uncertainty was greater when high changed to low (*t*_473_ = −7.78, *p <* 10^−13^, paired *t*-test), resulting in the faster learning rate (*t*_473_ = −7.91, *p <* 10^−13^, paired *t*-test). We also looked at the dynamics of the latent variables within blocks to see if they evolved on timescales relevant to behavior and task structure. While block lengths were prescribed to be 20–35 trials long, the block length experienced by the animal was often shorter (8.99 ± 2.78) due to the probabilities changing independently at each spout and the animals switching choices (which begins a new experienced block). We found that when entering a new block (from the animals’ perspective), expected uncertainty became lower in the high block relative to the medium block within approximately 5 trials (4.97 ± 1.51). The number of trials the model took to distinguish between reward probabilities in this way was less than the average experienced block lengths (Figure A2f,g, 9.03 ± 2.81, *t*_40_ = 8.67, *p <* 10^−10^, paired *t*-test). Thus, the updating rate of expected uncertainty allows for the calculation of expected uncertainty and detection of probability changes on timescales relevant to the task and behavior.

We also found evidence of meta-learning in the intra-trial lick behavior. Following no reward, mice consistently licked the chosen spout several times. We found that the number of licks was better explained by unexpected uncertainty from the meta-learning model than by RPE magnitude from the static learning model (Figure 2e). In other words, mice licked more when the no reward outcome was most unexpected.

## Serotonin neuron firing rates correlate with expected uncertainty

To quantify the link between serotonin neurons and meta-learning, we recorded action potentials from dorsal raphe serotonin neurons in mice performing the foraging task (66 neurons from 4 mice). To identify serotonin neurons, we expressed the light-gated ion channel, channelrhodopsin-2, under the control of the serotonin transporter promoter in *Slc6a4*-Cre (also known as *Sert*-Cre) mice (Figures 3a, A3a). We delivered light stimuli to the dorsal raphe to “tag” serotonin neurons at the end of each recording (Figures 3b, A3b,c). We calculated firing rates during the inter-trial intervals and compared the activity to the behavioral model variables. We found a significant relationship between firing rate and expected uncertainty in 53% (35 of 66) of serotonin neurons (Figure 3c-e; regression of inter-trial interval firing rates on expected uncertainty). When we regressed out slow, monotonic changes in firing rates and expected uncertainty over the course of the session, this relationship held (Figure A3d). In contrast, we did not find such prevalent relationships in a multivariate regression of firing rates on other latent model variables, such as relative value or RPE (Figure A3e).

**Figure 3.**
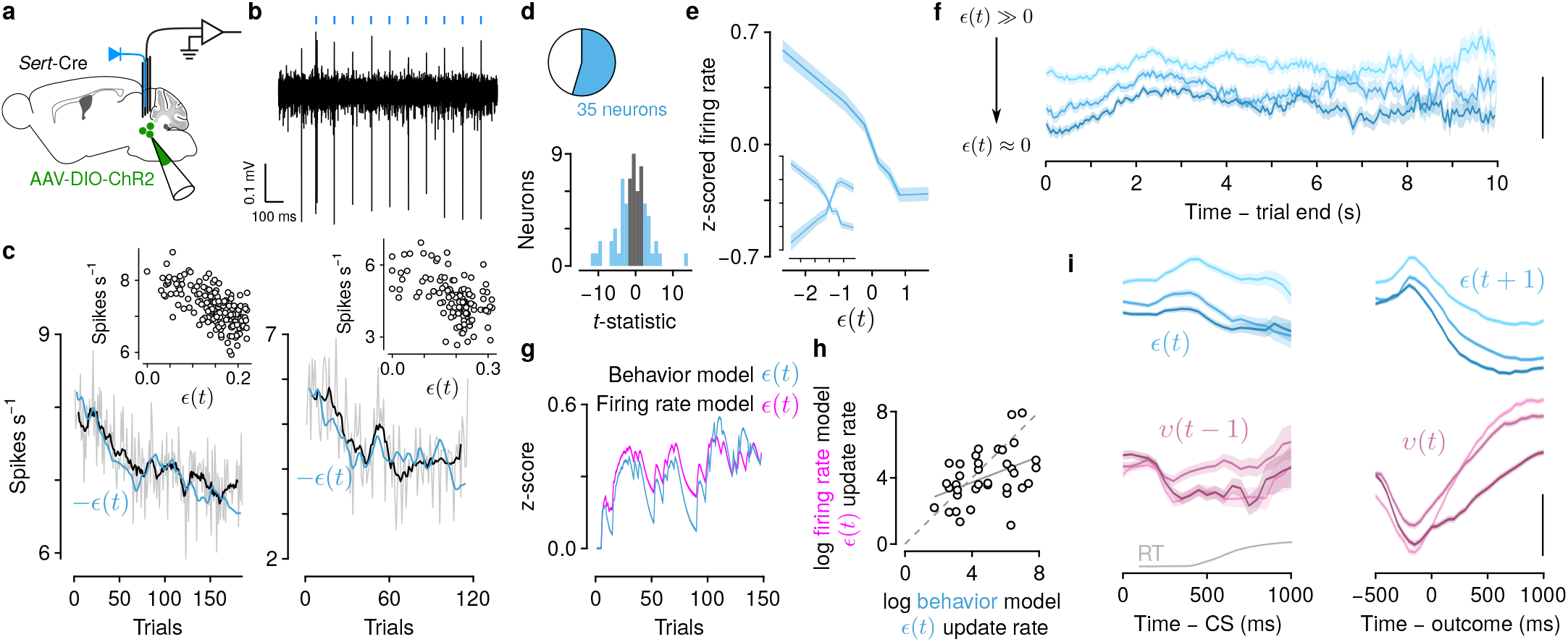
Serotonin neuron firing rates correlate with expected uncertainty on slow timescales and unexpected uncertainty on fast timescales. (a) Schematic of electrophysiological recording of identified serotonin neurons. (b) Example “tagging” of a serotonin neuron, using channelrhodopsin-2 stimulation. (c) Two example neurons showing negative correlations between inter-trial interval firing rates and expected uncertainty (*−ϵ*(*t*) is plotted). Insets show trial-by-trial relationships. (d) *t*-statistics across all neurons from a linear regression, modeling firing rates as a function of *ϵ*(*t*). Blue bars and slice indicate neurons with significant regression coefficients. (e) Population z-scored firing rates varied with *ϵ*(*t*). Inset shows population split by positive and negative correlations. “Sign-flipped” plot, combining across these neurons, was used for analysis in (f). (f) z-scored firing rates of serotonin neurons split by *ϵ*(*t*) tercile. Scale bar: 0.5 z-score. (g) Example dynamics of *ϵ*(*t*) estimated from behavior and neuronal firing rates. (h) Log-log plot of the expected uncertainty update rate (*ψ*) from each neuron’s firing rate model and the behavioral model derived from simultaneous choice behavior. (i) Within-trial dynamics of expected (*ϵ*(*t*), *ϵ*(*t* + 1), top row) and unexpected (*v*(*t −* 1), *v*(*t*), bottom row) uncertainty, aligned to go cue (CS, left column) and outcome (right column). Scale bar: 0.5 z-score. Gray curve: response time (RT) distribution (cut off at 1 s).

Remarkably, firing rates were stable within inter-trial intervals. Dividing expected uncertainty into terciles, we found that serotonin neuron firing rates were relatively constant as time elapsed within inter-trial intervals (Figure 3f; regression coefficient = 1.9 × 10^−6^ from a linear model of tercile difference on time in inter-trial interval). Because expected uncertainty evolved somewhat slowly as a function of RPE magnitude and the activity of neurons on this timescale (tens of seconds) fluctuated slowly as well, the two may be similarly autocorrelated (Elber-Dorozko and Loewenstein, 2018). To control for spurious correlations due to comparison of two autocorrelated variables, we first compared the actual neural data to simulated expected uncertainty terms (Figure A3f). We found stronger statistical relationships across the population with the actual expected uncertainty than with simulated values. Additionally, we simulated neural activity with quantitatively-matched autocorrelation functions to the real neurons and compared this activity to the actual expected uncertainty values. Again, we found stronger statistical relationships in the real data as opposed to the simulated data (Figure A3g-i).

To further examine the robustness of this relationship, we fit the meta-learning model to the inter-trial interval firing rates of neurons that had a significant correlation with expected uncertainty. The meta-learning algorithm was essentially the same as before, but we fit firing rates as a function of expected uncertainty as opposed to fitting choices as a function of relative action values. Here, we found that the updating rate for expected uncertainty from the firing rate model covaried with the same parameter from the choice model across sessions (Figure 3g,h). Additionally, how well the model fit to the firing rates was predicted by how well-correlated the firing rates were to the expected uncertainty variable from the behavioral model (Figure A3j). This result suggests that the neural and behavior data, independently, predict similar expected uncertainty dynamics.

## Serotonin neuron firing rates correlate with unexpected uncertainty at outcomes

How does the presence or absence of reward update the slowly-varying firing rates of serotonin neurons? According to the model, expected uncertainty changes as a function of unexpected uncertainty. In particular, the model thus predicts a firing rate change at the time of outcome that could be used to update expected uncertainty.

To test this, we calculated firing rates of serotonin neurons within trials, while mice made choices and received outcomes. We found that firing rate changes on fast timescales (hundreds of ms) correlated with expected uncertainty (*ϵ*(*t*)) throughout the period when mice received go cues and made choices (Figure 3i). These correlations persisted during the outcome (reward or no reward), as *ϵ*(*t*) updated to its next value (*ϵ*(*t* + 1)). In contrast, firing rates correlated with unexpected uncertainty (*v*(*t*)) primarily during the outcome, but not during the go cue (*v*(*t* − 1); Figure 3i). Thus, brief firing rate changes in serotonin neurons could be integrated to produce more slowly-varying changes. In this computation, firing rates may be interpreted as encoding two forms of uncertainty, one slowly varying (*ϵ*), one more transient (*v*).

## Serotonin neuron firing rates correlate with uncertainty in a Pavlovian task

Based on the results from the first experiment, we made two predictions. First, we hypothesized that correlations between serotonin neuron activity and expected uncertainty generalize to other behavioral tasks. To test this, we trained 9 mice on a Pavlovian version of the task in which an odor cue predicted probabilistic reward after a 1-s delay (Figure 4a). The probability of reward changed in blocks within each session (Figure 4b). This task required no choice to be made. Rather, mice simply licked toward a single water-delivery spout in anticipation of a possible reward.

**Figure 4.**
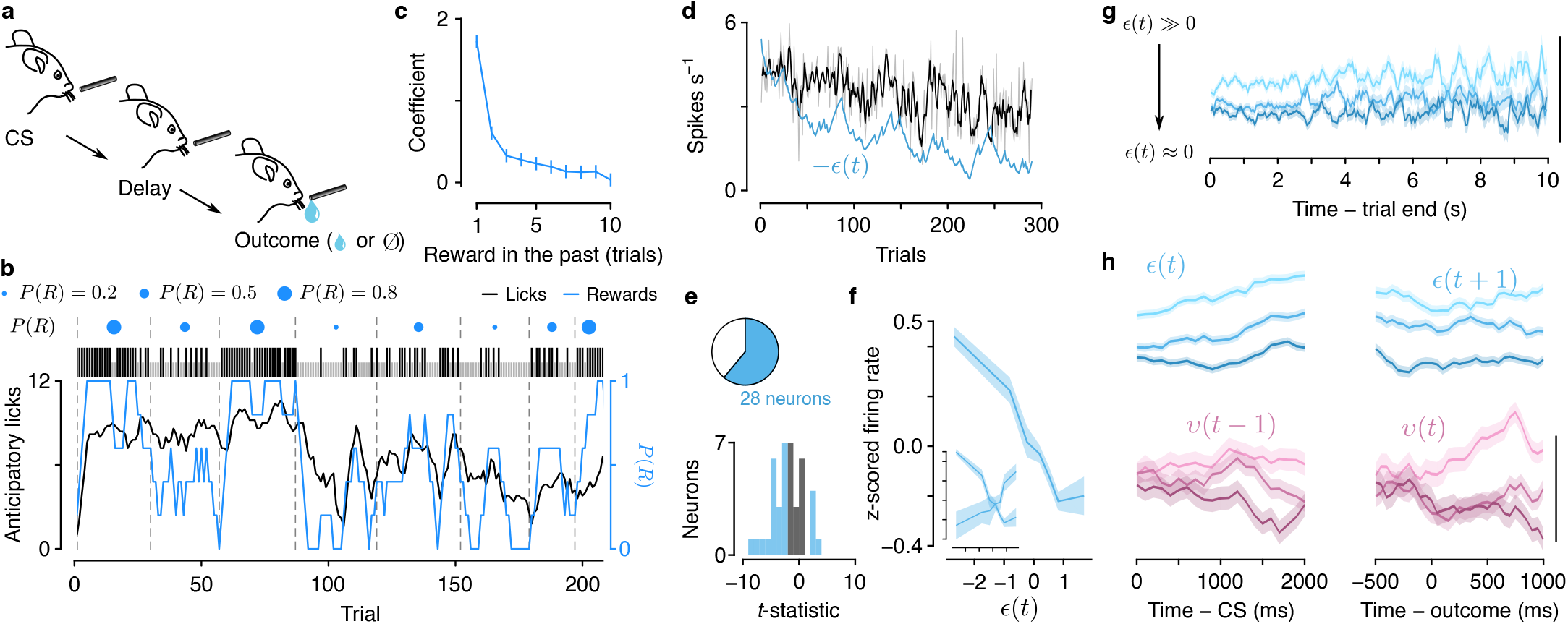
Serotonin neuron firing rates correlate with expected and unexpected uncertainty in a dynamic Pavlovian task. (a) Schematic of Pavlovian task in which the probability of reward (*P* (*R*)) varied over trials. (b) Example behavior showing anticipatory licking, in the delay before outcome, as *P* (*R*) varied. Black ticks: rewarded trials. Gray ticks: unrewarded trials. (c) Linear regression coefficients of licking rate on reward history. (d) Example serotonin neuron showing a negative correlation between inter-trial interval firing rates and expected uncertainty (*−ϵ*(*t*) is plotted). (e) *t*-statistic from linear regression, modeling firing rate as a function of *ϵ*(*t*) as in Figure 3d. (f) Population “tuning curves,” as in Figure 3e. (g) Stable firing rates within inter-trial intervals, as in Figure 3f. Scale bar: 0.5 z-score. (h) Expected and unexpected uncertainty z-scored firing rates, as in Figure 3i. Scale bar: 0.5 z-score.

The number of anticipatory licks during the delay between cue and outcome (presence or absence of reward) reflected recent reward history (Figure 4c). To estimate ongoing expected uncertainty in this task, we modified the meta-learning model to generate anticipatory licks as a function of the value of the cue (Figure A4a). While the model was capable of explaining behavior, interestingly, we found no clear behavioral evidence of variable learning rates (Figure A4b,c). However, recordings of dorsal raphe serotonin neurons from mice behaving in this task revealed that the activity of these neurons correlated with expected uncertainty at similar rates to those recorded in the dynamic foraging task (Figures 4d-f, A4d-f; 68%, 28 of 41 neurons from 5 mice). Similar to observations in the foraging task, neurons in the Pavlovian task showed stable firing rates within inter-trial intervals (Figure 4g; regression coefficient = −3.1 × 10^−6^ from a linear model of tercile difference on time in inter-trial interval). Serotonin neuron firing rates also correlated with expected uncertainty throughout its update interval, and with unexpected uncertainty at the time of the outcome (Figure 4h). Thus, the nervous system may maintain running estimates of two forms of uncertainty that generalizes across behavioral tasks.

## Serotonin neuron inhibition disrupts meta-learning

In our second prediction from the dynamic foraging experiment, we asked whether inactivating serotonin neurons rendered mice unable to adjust learning rates. The meta-learning model makes specific predictions about the role of uncertainty in learning. To test the predictions of the model under the hypothesis that serotonin neurons encode expected uncertainty, we expressed an inhibitory designer receptor exclusively activated by designer drugs (DREADD) conjugated to a fluorophore (hM4Di-mCherry) in dorsal raphe serotonin neurons (Figure 5a). *Sert*-Cre mice received injections of a Cre-dependent virus containing the receptor (AAV5-hSyn-DIO-hM4D(Gi)-mCherry, *n* = 3 mice) into the dorsal raphe. Control *Sert*-Cre mice were injected with the same virus containing only the fluorophore (*n* = 4 mice). On consecutive days, mice received an injection of vehicle (0.5% DMSO in 0.9% saline), the DREADD ligand agonist 21 (3 mg kg^−1^ in vehicle; Chen et al., 2015; Thompson et al., 2018), or no injection. Because the simultaneous changes of reward probabilities were rare, we modified the task to include them with slightly higher frequency.

**Figure 5.**
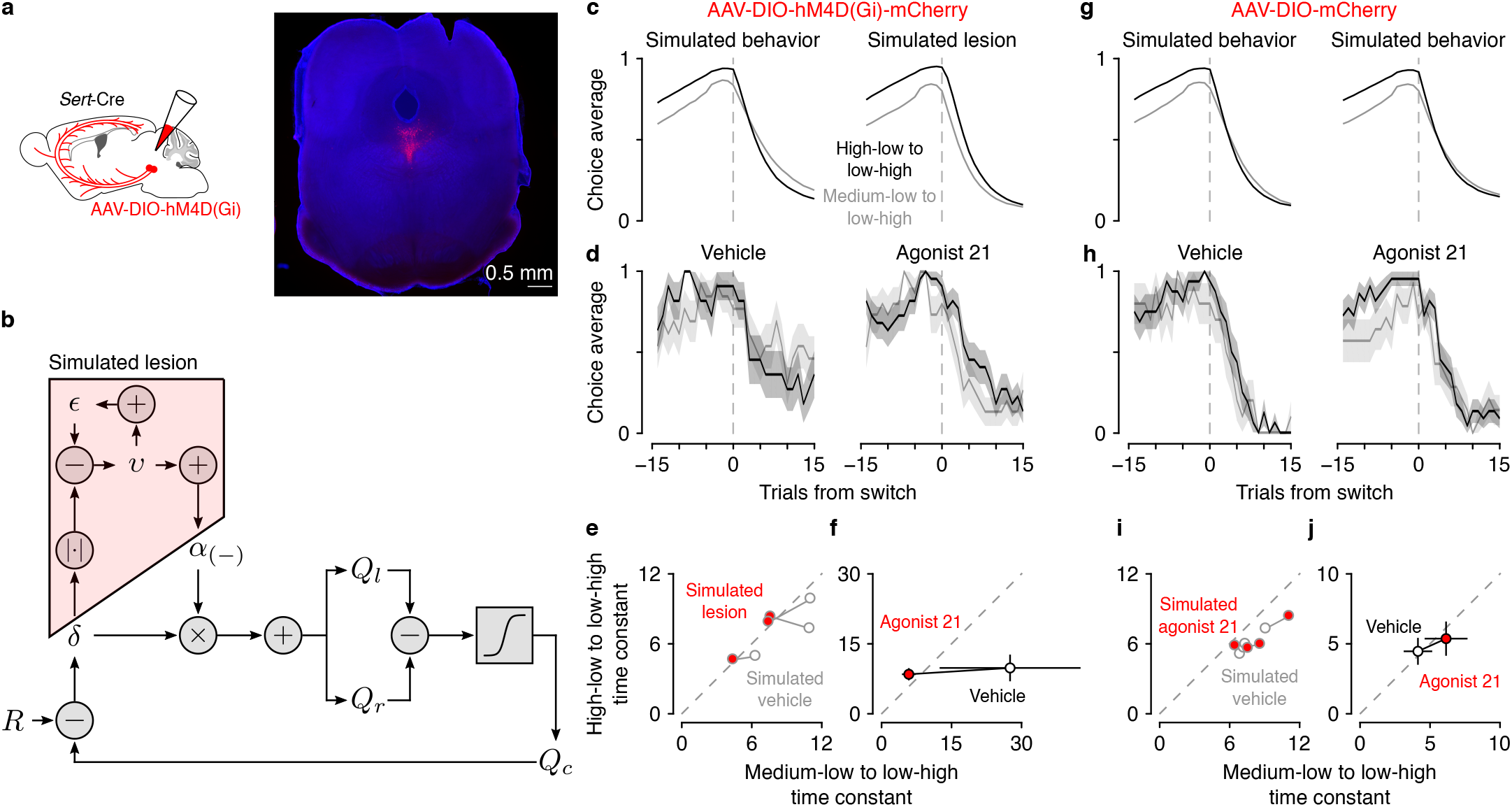
Serotonin neuron inhibition disrupts meta-learning. (a) Schematic of experiment to reversibly inactivate serotonin neurons and representative expression of hM4Di-mCherry in dorsal raphe serotonin neurons. (b) Schematic of simulated lesion, in which models were fit to mouse behavior, and then meta-learning variables (i.e., *ϵ* and *v*) were set to zero. (c) Simulated behavior with meta-learning intact, fit to vehicle behavior (left) and simulated lesion (right). (d) Mouse behavior with vehicle injections (control experiment) and drug (agonist 21). Lines are mean choice probability and shading is Bernoulli S.E.M. (e) Exponential time constants for transitions from simulated behavior and simulated lesions. (f) Time constants from mice (with 95% C.I.). (g) Simulated behavior from mice expressing mCherry in serotonin neurons with vehicle (left) and agonist 21 (right) injections. (h) Mouse behavior with vehicle injections (control experiment) and drug (agonist 21). (i) Simulation time constants from fluorophore-control mice. (j) Time constants from fluorophore-control mice (with 95% C.I.).

To quantify the change in behavior predicted by the model, we first fit the model to mouse behavior on vehicle injection days and used those parameters to simulate behavior. We then simulated a lesion by fixing expected and unexpected uncertainty to 0 (essentially fixing the negative RPE learning rate to its median value) and simulated behavior again (Figure 5b). The simulated lesion diminished the differences in transition speed between the pre-transition reward conditions (Figure 5c,e).

On days with agonist 21 injections, mice expressing hM4Di in serotonin neurons demonstrated changes in learning at transitions (Figure 5d,f; effects of trial from transition *F*_1,58_ = 44.3, *p <* 10^−7^, drug condition *F*_1,58_ = 41.5, *p <* 10^−7^, trial × transition type interaction *F*_1,58_ = 6.15, *p* = 0.016, and transition type × drug condition interaction *F*_1,58_ = 21.2, *p <* 10^−4^, linear mixed effects model) matching the predictions of the simulated lesion model (Figure 5c,e; effects of trial from transition *F*_1,186_ = 180.4, *p <* 10^−28^, drug condition *F*_1,186_ = 18.1, *p <* 10^−4^, trial × transition type interaction *F*_1,186_ = 7.51, *p* = 0.0067, and transition type × drug condition interaction *F*_1,186_ = 8.28, *p* = 0.0045). The same experiment in mice expressing a fluorophore alone in serotonin neurons showed no effect of agonist 21 (Figure 5h,j; effect of trial from transition *F*_1,58_ = 95.8, *p <* 10^−13^ and effect of transition type *F*_1,58_ = 5.55, *p* = 0.022), consistent with simulations from the meta-learning model fit separately to vehicle and agonist 21 behavior (Figure 5g,i; effect of trial from transition *F*_1,250_ = 483, *p <* 10^−59^ and effect of trial type × transition type interaction *F*_1,58_ = 7.02, *p* = 0.0086). Serotonin neuron inhibition did not slow response times (Figure A5b), change how outcomes drove response times (Figure A5c), nor cause mice to lick during inter-trial intervals. Thus, the observed effects of reversible inhibition are consistent with a role for serotonin neurons signaling uncertainty.

## Discussion

To behave flexibly in dynamic environments, learning rates should vary according to the statistics of those environments (Dayan et al., 2000; Doya, 2002; Kakade and Dayan, 2002; Yu and Dayan, 2005; Behrens et al., 2007). Our model captures differences in learning by estimating expected uncertainty: a moving average of unsigned prediction errors that tracks variability in the outcomes of actions. This quantity is used to modulate learning rate by determining how unexpected an outcome is relative to that expected uncertainty. When outcomes are probabilistic but stable, expected uncertainty also slows learning. The model captured observed changes in learning rates that could not be reproduced with an RL model that uses static learning rates. The activity of the majority of identified serotonin neurons correlated with the expected uncertainty variable from the model when fit to dynamic foraging behavior. This relationship held in a different behavioral context, with similar fractions of serotonin neurons tracking expected uncertainty in a dynamic Pavlovian task. During dynamic foraging, chemogenetic inhibition of serotonin neurons caused changes in choice behavior that were consistent with the changes in learning predicted by removing meta-learning from the model.

While serotonin neuron firing rates change on multiple timescales (Cohen et al., 2015), the observed changes that correlated with expected uncertainty occurred over relatively long periods of time. How rapidly RPE magnitudes are integrated tracks variability in outcomes on a timescale relevant to experienced block lengths. Consequently, deviations from expected uncertainty reliably indicate changes in reward probabilities. In addition to the computational relevance of activity on this timescale, serotonin neuron firing rate changes may be optimized for the nervous system to implement these computational goals. Slow changes in serotonin neuron activity could enable gating or gain control mechanisms (Shimegi et al., 2016; Azimi et al., 2020), bidirectional modulation of relevant inputs and outputs (Avesar and Gulledge, 2012; Stephens et al., 2014, 2018), or other previously-observed circuit mechanisms that modulate how new information is incorporated (Marder, 2012) to drive flexible behavior.

We also observed changes in serotonin neuron activity on shorter timescales that correlated with both expected and unexpected uncertainty. The timing of these brief signals may be important to update the slower dynamics correlated with expected uncertainty, as predicted from the model (i.e., *ϵ* essentially integrates *v*). Alternatively, serotonin neurons could “multiplex” across timescales, whereby brief changes in firing rates may have different downstream functions than slower changes.

Several conceptualizations of expected uncertainty have been proposed with different consequences for learning and exploratory behavior (Soltani and Izquierdo, 2019). For example, there can be uncertainty about a specific causal relationship between events in the environment, or between a specific action and the environment. There is evidence that the activity of norepinephrine and acetylcholine neurons may be related to these types of uncertainty (Yu and Dayan, 2005; Hangya et al., 2015; Zhang et al., 2019). It should be noted that both norepinephrine neurons in the locus coeruleus (Szabo and Blier, 2001) and acetylcholine neurons in the basal forebrain (Bengtson et al., 2004) receive functional input from dorsal raphe serotonin neurons.

Here, we studied a more general form of expected uncertainty that tracks variability in outcomes regardless of the specific action taken. This type of uncertainty may apply to learned rules or separately, states in a model-based framework (Bach and Dolan, 2012). It may also be conceptually related to the level of commitment to a belief, which can scale learning in models that learn by minimizing surprise (Payzan-LeNestour and Bossaerts, 2011; Payzan-LeNestour et al., 2013; Faraji et al., 2018). In these ways, our model may approximate inference or change detection in certain behavioral contexts. Our notion of expected uncertainty is also related to reward variance, risk, or outcome uncertainty (Preuschoff et al., 2006; Preuschoff and Bossaerts, 2007; Bach and Dolan, 2012; Monosov, 2020), but with respect to an entire behavioral policy as opposed to a specific action.

Unexpected uncertainty has also been previously defined in numerous ways. In our model, the negative RPE learning rate is a function of recent deviations from expected uncertainty and thus may be most related to a subjective estimate of environmental volatility. This interpretation is consistent with learning rates increasing as a function of increasing volatility (Behrens et al., 2007). An estimate of volatility may also reflect the surprise that results from the violation of a belief (Payzan-LeNestour and Bossaerts, 2011; Payzan-LeNestour et al., 2013; Faraji et al., 2018). Our observation that brief changes in serotonin neuron firing rates at the time of outcome correlated with unexpected uncertainty is also consistent with previous work showing that serotonin neuron activity correlated with “surprise” when cue-outcome relationships were violated (Matias et al., 2017).

We did not find any evidence in the dynamic Pavlovian behavior that distinguished meta-learning from static learning. It may be that differences in these models are not observable in this behavior. Also, the dynamic foraging task engages regions of the brain that are not necessary for the dynamic Pavlovian task (Bari et al., 2019). Consequently, uncertainty may be incorporated in other ways to drive behavior. Alternatively, the brain may keep track of statistics of the environment that are not always used in behavior.

In the meta-learning RL model as we have formulated it, only the negative RPE learning rate is subject to meta-learning. This is an empirical finding and one that may be a consequence of the structure of the task. For example, the reward statistics might result in a saturation of learning from rewards such that its modulation is unnecessary. Asymmetries in the task structure (the absence of trials in which *P* (*R*) = 0.1 for both spouts) and mouse preference (mice regularly exploited the *P* (*R*) = 0.5 spout) also result in rewards carrying more information about which spout is “good enough” (*P* (*R*) = 0.9 or *P* (*R*) = 0.5). Another possibility, not mutually exclusive with the first, is that learning about rewards and lack thereof could be asymmetric. This asymmetry could result from ambiguity in the non-occurrence of the expected outcome, differences in the magnitude of values of each outcome, or separate learning mechanisms entirely. Similarly, because outcomes are binary in our tasks, learning from negative and positive RPEs could be asymmetric. Alternatively, as described above, this parameterization might just better approximate a more complex cognitive process (e.g., inference) in this specific behavioral context.

Our findings and conceptual framework, including the effect on learning from worse-than-expected outcomes, are consistent with previous observations and manipulations of serotonin neuron activity. In a Pavlovian reversal task, changes in cue-outcome mappings elicited responses from populations of serotonin neurons that decayed as mice adapted their behavior to the new mapping (Matias et al., 2017). Chemogenetic inhibition of serotonin neurons in this task impaired behavioral adaptation to a cue that predicted reward prior to the reversal but not after. The manipulation did not affect behavior changes in response to the opposite reversal. In reversal tasks in which action-outcome contingencies were switched, lesions or pharmacological manipulations of serotonin neurons also resulted in impairments of adaptive behavior at the time of reversal (Clarke et al., 2004, 2007; Boulougouris and Robbins, 2010; Bari et al., 2010; Brigman et al., 2010). Specifically, lesioned animals continued to make the previously-rewarded action. These findings are consistent with a role for serotonin neuron activity in tracking expected uncertainty and driving learning from worse-than-expected outcomes. More recent work demonstrated that serotonin neuron activation increased the learning rate after longer intervals between outcomes, but that learning after shorter intervals was already effectively saturated (i.e., win-stay, lose-shift; Iigaya et al., 2018). An intriguing possibility is that serotonin neurons mediate the contributions of faster, working-memory-based learning, and slower, plasticity-dependent learning that may map onto model-based and model-free learning, respectively (Balleine and Dickinson, 1998; Daw et al., 2005; Lee et al., 2014; Miller et al., 2017; Iigaya et al., 2018).

A number of studies have also examined the role of serotonin neuron activity in patience and persistence for rewards (Miyazaki et al., 2011, 2014; Fonseca et al., 2015; Lottem et al., 2018). These studies demonstrated that activating serotonin neurons increased waiting times for or active seeking of reward. In all cases, animals can be thought of as learning from lack of rewards at each point in time. Under the proposed meta-learning framework, increasing expected uncertainty would slow this learning, resulting in prolonging waiting times or enhancing persistence.

What are the postsynaptic consequences of slow changes in serotonin release? Target regions involved in learning and decision making, like the prefrontal cortex, ventral tegmental area, and striatum, express a diverse range of serotonin receptors capable of converting a global signal into local changes in circuit dynamics. The activity in these regions also correlates with latent decision variables that update with each experience (Schultz et al., 1997; Samejima et al., 2005; Lau and Glimcher, 2008; Massi et al., 2018; Wang et al., 2018; Bari et al., 2019), providing a potential substrate through which serotonin could modulate learning. For example, the gain of RPE signals produced by dopamine neurons in the ventral tegmental area is modulated by the variance of reward value (Fiorillo et al., 2003; Tobler et al., 2005).

What is the presynaptic origin of uncertainty computation in serotonin neurons? Synaptic inputs from the prefrontal cortex (Geddes et al., 2016) may provide information about decision variables used in this task (Bari et al., 2019). Local circuit mechanisms in the dorsal raphe (Geddes et al., 2016; Zhou et al., 2017) and long-lasting conductances in serotonin neurons (Haj-Dahmane et al., 1991; Penington et al., 1993; Brown et al., 2002; Andrade et al., 2015; Gantz et al., 2020) likely contribute to the persistence of these representations.

Learning is dynamic. Flexible decision making requires using recent experience to adjust learning rates adaptively. The observed foraging behavior demonstrates that learning is not a static process, but a dynamic one. The meta-learning RL model provides a potential mechanism by which recent experience modulates learning adaptively, and reveals a quantitative link between serotonin neuron activity and flexible behavior.

## Acknowledgments

We thank Terry Shelley for machining, Drs. Daeyeol Lee, David Linden, Daniel O’Connor, Marshall Hussain Shuler, and the lab of Reza Shadmehr for comments, and Dr. Michael Betancourt for advice on model fitting. This work was supported by Klingenstein-Simons, MQ, NARSAD, Whitehall, R01DA042038, and R01NS104834 (J.Y.C.), and P30NS050274.

## Author contributions

C.D.G. and B.A.B. collected data. C.D.G., B.A.B., and J.Y.C. designed experiments, analyzed data, and wrote the paper.

## Methods

### Animals and surgery

We used 53 male and female mice, backcrossed with C57BL/6J and heterozygous for Cre recombinase under the control of the serotonin transporter gene (Slc6a4^*tm*1(*cre*)*Xz*^, The Jackson Laboratory, 014554; Zhuang et al., 2005). 4 mice were used for electrophysiological recordings in the dynamic foraging task (4 male), 5 mice were used for electrophysiological recordings in the dynamic Pavlovian task (1 female, 4 male), 37 mice (18 female, 19 male) were used for additional behavior in the dynamic foraging task, 4 mice (4 male) were used for additional behavior in the dynamic Pavlovian task, and 7 mice (3 female, 4 male) were used for the chemogenetic experiments. Surgery was performed on mice between the ages of 4–8 weeks, under isoflurane anesthesia (1.0–1.5% in O_2_) and in aseptic condtions. During all surgeries, custom-made titanium headplates were surgically attached to the skull using dental adhesive (C&B-Metabond, Parkell). After the surgeries, analgesia (ketoprofen, 5 mg kg^−1^ and buprenorphine, 0.05–0.1 mg kg^−1^) was administered to minimize pain and aid recovery.

For electrophysiological experiments, we implanted a custom microdrive targeting dorsal raphe using a 16° posterior angle, entering through a craniotomy at 5.55 mm posterior to bregma and aligned to the midline.

For all experiments, mice were given at least one week to recover prior to water restriction. During water restriction, mice had free access to food and were monitored daily in order to maintain 80% of their baseline body weight. All mice were housed in reverse light cycle (12h dark/12h light, dark from 08:00–20:00) and all experiments were conducted during the dark cycle between 10:00 and 18:00. All surgical and experimental procedures were in accordance with the *National Institutes of Health Guide for the Care and Use of Laboratory Animals* and approved by the Johns Hopkins University Animal Care and Use Committee.

### Behavioral task

Before training on the tasks, water-restricted mice were habituated to head fixation for 1–3 d with free access to water from the provided spouts (two 21 ga stainless steel tubes separated by 4 mm) placed in front of the 38.1 mm acrylic tube in which the mice rested. The spouts were mounted on a micromanipulator (DT12XYZ, Thorlabs) with a custom digital rotary encoder system to reliably determine the position of the lick spouts in XYZ space with 5–10 *μ*m resolution (Bari et al., 2019). Each spout was attached to a solenoid (ROB-11015, Sparkfun) to enable retraction (see Behavioral tasks: dynamic foraging). The odors used for the cues (p-cymene and (−)-carvone) were dissolved in mineral oil at 1:10 dilution (30 *μ*l) and absorbed in filter paper housed in syringe adapters (Whatman, 2.7 *μ*m pore size). The adapters were connected to a custom-made olfactometer (Cohen et al., 2012) that diluted odorized air with filtered air by 1:10 to produce a 1.0 L min^−1^ flow rate. The same flow rate was maintained outside of the cue period so that flow rate was constant throughout the task.

Licks were detected by charging a capacitor (MPR121QR2, Freescale) or using a custom circuit (Janelia Research Campus 2019-053). Task events were controlled and recorded using custom code (Arduino) written for a microcontroller (ATmega16U2 or ATmega328). Water rewards were 2–4 *μ*l, adjusted for each mouse to maximize the number of trials completed per session and to keep sessions around 60 minutes. Solenoids (LHDA1233115H, The Lee Co) were calibrated to release the desired volume of water and were mounted on the outside of the dark, sound-attenuated chamber used for behavior tasks. White noise (2–60 kHz, Sweetwater Lynx L22 sound card, Rotel RB-930AX two-channel power amplifier, and Pettersson L60 Ultrasound Speaker), was played inside the chamber to block any ambient noise.

### Behavioral tasks: dynamic foraging

During the 1–3 days of habituation, mice were trained to lick both spouts to receive water. Water delivery was contingent upon a lick to the correct spout at any time. Reward probabilities were chosen from the set {0, 1} and reversed every 20 trials.

In the second stage of training (5–12 d), the trial structure with odor presentation was introduced. Each trial began with the 0.5 s delivery of either an odor “go cue” (*P* = 0.95) or an odor “no-go cue” (*P* = 0.05). Following the go cue, mice could lick either the left or the right spout. If a lick was made during a 1.5 s response window, reward was delivered probabilistically from the chosen spout. The unchosen spout was retracted at the time of the tongue contacting the other spout so that mice would not try to sample both spouts within a trial. The unchosen spout was replaced 2.5 s after cue onset. Following a no-go cue, any lick responses were neither rewarded nor punished. Reward probabilities during this stage were chosen from the set {0, 1} and reversed every 20–35 trials. During this period of training only, water was occasionally manually delivered to encourage learning of the response window and appropriate switching behavior. Reward probabilities were then changed to {0.1, 0.9} for 1–2 days of training prior to introducing the final stage of the task. Rewards were never “baited,” as in previous versions of the task (Sugrue et al., 2004; Lau and Glimcher, 2005; Tsutsui et al., 2016; Bari et al., 2019). We did not penalize switching with a “changeover delay.” If a directional lick bias was observed in one session, the lick spouts were moved horizontally 50–300 *μ*m in the opposite direction prior to the following session.

After the 1.5 s response window, inter-trial intervals were generated as draws from an exponential distribution with a rate parameter of 0.3 and a maximum of 30 s. This distribution results in a flat hazard rate for inter-trial intervals such that the probability of the next trial did not increase over the duration of the inter-trial interval (Luce, 1986). Inter-trial intervals (go-cue on to go-cue on) were 7.45 s on average (range 2.5–32.5 s). As in previous studies, mice made a leftward or rightward choice in greater than 99% of trials (Bari et al., 2019). Mice completed 280 ± 66.6 trials per session (range 79–655 trials).

In the final stage of the task, the reward probabilities assigned to each lick spout were drawn pseudorandomly from the set {0.1, 0.5, 0.9} in all the mice from the behavior experiments (*n* = 48), all the mice from the DREADDs experiments (*n* = 7), and half of the mice from the electrophysiology experiments (*n* = 2). The other half of mice from the electrophysiology experiments (*n* = 2) were run on a version of the task with probabilities drawn from the set {0.1, 0.4, 0.7}. The probabilities were assigned to each spout individually with block lengths drawn from a uniform distribution of 20–35 trials. To stagger the blocks of probability assignment for each spout, the block length for one spout in the first block of each session was drawn from a uniform distribution of 6–21 trials. For each spout, probability assignments could not be repeated across consecutive blocks. To maintain task engagement, reward probabilities of 0.1 could not be simultaneously assigned to both spouts. If one spout was assigned a reward probability greater than or equal to the reward probability of the other spout for 3 consecutive blocks, the probability of that spout was set to 0.1 to encourage switching behavior and limit the creation of a direction bias. If a mouse perseverated on a spout with reward probability of 0.1 for 4 consecutive trials, 4 trials were added to the length of both blocks. This procedure was implemented to keep mice from choosing one spout until the reward probability became high again.

To minimize spontaneous licking, we enforced a 1 s no-lick window prior to odor delivery. Licks within this window were punished with a new randomly-generated inter-trial interval, followed by a 2.5 s no-lick window. Implementing this window significantly reduced spontaneous licking throughout the entirety of behavioral experiments.

### Behavioral tasks: dynamic Pavlovian

On each trial either an odor “CS+” (*P* = 0.95) or an odor “CS−” (*P* = 0.05) was delivered for 1 s followed by a delay of 1 s. CS+ predicted probabilistic reward delivery, whereas CS− predicted nothing. Mice were allowed 3 s to consume the water, after which any remaining reward was removed by a vacuum. Each trial was followed by an inter-trial interval, drawn from the same distribution as in the dynamic foraging task. The time between trials (CS on to CS on) was 9.34 s on average (range 6–36 s).

The reward probability assigned to CS+ was drawn pseudorandomly from the set {0.2, 0.5, 0.8} or, in separate sessions, alternated between the probabilities in the set {0.2, 0.8}. The probability changed every 20–70 trials (uniform distribution). The CS+ probability of the first block of every session was 0.8.

### Electrophysiology

We recorded extracellular signals from neurons at 32 or 30 kHz using a Digital Lynx 4SX (Neuralynx Inc,) or Intan Technologies RHD2000 system (with RHD2132 headstage), respectively. The recording systems were connected to 8–16 implanted tetrodes (32–64 channels, nichrome wire, PX000004, Sandvik) fed through 39 ga polyimide guide tubes that could be advanced with the turn of a screw on a custom, 3D-printed microdrive. The impedances of each wire in the tetrodes were reduced to 200–300 kΩ by gold plating. The tetrodes were wrapped around a 200 *μ*m optic fiber used for optogenetic identification. After each recording session, the tetrode-optic-fiber bundle was driven down 75 *μ*m. The median signal was subtracted from raw recording traces across channels and bandpass-filtered between 0.3–6 kHz using custom MATLAB software. To detect peaks, the bandpass-filterd signal, *x*, was thresholded at 4*σ*_*n*_ where 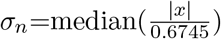 (Quiroga et al., 2004). Detected peaks were sorted into individual unit clusters offline (Spikesort 3D, Neuralynx Inc.) using waveform energy, peak waveform amplitude, minimum waveform trough, and waveform prinipal component analysis. We used two metrics of isolation quality as inclusion criteria: L-ratio (*<* 0.05) (Schmitzer-Torbert et al., 2005) and fraction of interspike interval violations (*<* 0.1% interspike intervals *<* 2 ms).

Individual neurons were determined to be optogenetically-identified if they responded to brief pulses (10 ms) of laser stimulation (473 nm wavelength) with short latency, small latency variability, and high probability of response across trains of stimulation (10 trains of 10 pulses delivered at 10 Hz). We used an unsupervised k-means clustering algorithm to cluster all neurons based on these features. The elbow method and Calinkski-Harabasz criterion were used to determine that the optimal number of clusters was 4. Members of the cluster (66 neurons) with the highest mean probability of response, shortest mean latency, and smallest mean latency standard deviation were considered as identified. The responses of individual neurons were manually inspected to ensure light responsivity. In addition to the presence of identified serotonin neurons, targeting of dorsal raphe was confirmed by performing electrolytic lesions of the tissue (20 s of 20 *μ*A direct current across two wires of the same tetrode) and examining the tissue after perfusion.

### Viral injections

To express channelrhodopsin-2 (ChR2), hM4di, or mCherry in dorsal raphe serotonin neurons, we pressure-injected 810 nl of rAAV5-EF1a-DIO-hChR2(H134R)-EYFP (3 × 10^13^ GC ml^−1^), pAAV-hSyn-DIO-hM4D(Gi)-mCherry (1.2 × 10^13^ GC ml^−1^), or pAAV-hSyn-DIO-mCherry (1.0 × 10^13^ GC ml^−1^) into the dorsal raphe of *Sert*-Cre mice at a rate of 1 nl/s (MMO-220A, Narishige). pAAV-hSyn-DIO-hM4D(Gi)-mCherry was a gift from Bryan Roth (Addgene viral prep 44362-AAV5). We made three injections of 270 nl at the following coordinates: {4.63, 4.57, 4.50} mm posterior of bregma, {0.00, 0.00, 0.00} mm lateral from the midline, and {2.80, 3.00, 3.25} mm ventral to the brain surface. The pipette was inserted through a craniotomy at −5.55 mm posterior to bregma and aligned to midline, using a 16° posterior angle. Before the first injection, the pipette was left at the most ventral coordinate for 10 minutes. After each injection, the pipette was withdrawn 50 *μ*m and left in place for 5 min. The craniotomy after a hM4Di or mCherry injection was covered with silicone elastomer (Kwik-Cast, WPI) and dental cement. For electrophysiology experiments with rAAV5-EF1a-DIO-hChR2(H134R)-EYFP injections, the microdrive was implanted through the same craniotomy.

### Inactivation of serotonin neurons

Four mice were injected with pAAV-hSyn-DIO-hM4D(Gi)-mCherry and 4 mice were injected with pAAV-hSyn-DIO-mCherry as a control. One of the hM4D mice failed to perform the task and so was excluded. After training mice, we injected either 3.0 mg kg^−1^ agonist 21 (Tocris) dissolved in 0.5% DMSO/saline or an equivalent volume of vehicle (0.5% DMSO/saline alone) I.P. on alternating days (5 sessions per injection type per mouse).

### Data analysis

All analyses were performed with MATLAB (Mathworks) and R. All data are presented as mean ± S.D. unless reported otherwise. All statistical tests were two-sided. In Figure 2d, the probability that the time constants from the actual behavior belonged to the distribution of simulated behavior time constants was calculated by finding the Mahalanobis distance of the former from the latter, calculating the cumulative density function of the chi-square distribution at that distance, and subtracting it from 1. For all analyses, no-go (dynamic foraging) and CS− (dynamic Pavlovian) cues were ignored and treated as part of the inter-trial interval.

### Data analysis: descriptive models of behavior

We fit logistic regression models to predict choice as a function of outcome history for each mouse using the model

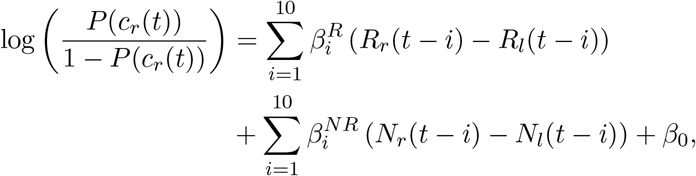

where *c*_*r*_(*t*) = 1 for a right choice and 0 for a left choice, *R* = 1 for a rewarded choice and 0 for an unrewarded choice, and *N* = 1 for an unrewarded choice and 0 for a rewarded choice. To predict response times (RT), we first z-scored the lick latencies by spout, to correct for differences due to relative spout placement and bias. Then, for each animal we fit the model

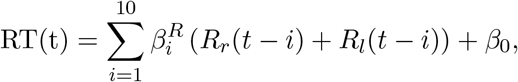

including a variable for trial number. We fit exponentials with the equation 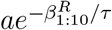 to the regression coefficients, averaged across animals, from the choice and response time models.

### Data analysis: generative model of behavior with static learning

We applied a generative RL model of behavior in the foraging task with static learning rates (Daw et al., 2006; Bari et al., 2019). This RL model estimates action values (*Q*_*l*_(*t*) and *Q*_*r*_(*t*)) on each trial to generate choices. Choices are described by a random variable, *c*(*t*), corresponding to left or right choice, *c*(*t*) ∈ {*l, r*}. The value of a choice is updated as a function of the RPE, and the rate at which this learning occurs is controlled by the learning rate parameter *α*. Because we observed asymmetric learning from rewards and no rewards (Figure 1c), consistent with previous reports (Bari et al., 2019), we included separate learning rates for the different outcomes. For example, if the left spout was chosen, then

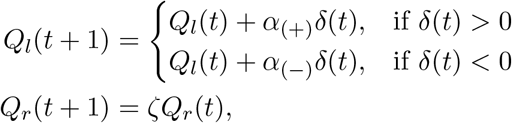

where *δ*(*t*) = *R*(*t*) − *Q*_*l*_(*t*) and *ζ* represents the forgetting rate parameter. The forgetting rate captures the increasing uncertainty about the value of the unchosen spout.

The *Q*-values are used to generate choice probabilities through a softmax decision function (Daw et al., 2006):

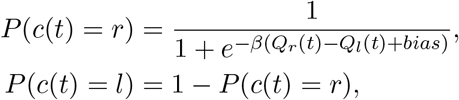

where *β*, the “inverse temperature” parameter, controls the steepness of the sigmoidal function. In other words, *β* controls the level of exploration versus exploitation with respect to the relative action values.

### Data analysis: generative model of behavior with meta-learning

We observed mouse behavior that the static learning model failed to capture and that suggested that learning rate was not constant over time. Thus, we added a component to the model that modulates RPE magnitude and *α*_(−)_ (“meta-learning”). Because learning should be slow in stable but variable environments, expected uncertainty scaled RPEs, such that learning is decreased when expected uncertainty is high. If the left spout was chosen, the values of actions were updated according to

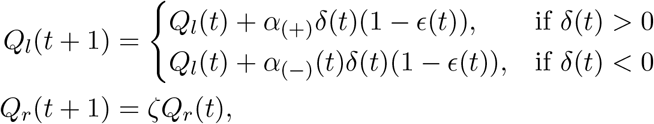

where *ϵ* is an evolving estimate of expected uncertainty calculated from the history of unsigned RPEs:

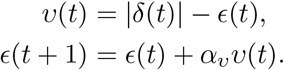

The rate of RPE magnitude integration is controlled by *α*_*v*_. Deviations from the expected uncertainty are captured by unexpected uncertainty, *v*, and may indicate that a change has occurred in the environment. Changes in the environment should drive learning to adapt behavior to new contingencies so *α*_(−)_ varies as a function of how surprising recent outcomes are:

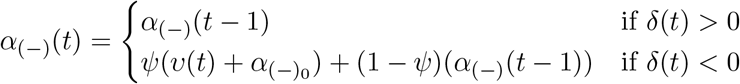

where 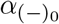 is the baseline learning rate from no reward and *ψ* controls how quickly unexpected uncertainty is integrated to update *α*_(−)_. As it is formulated, *α*_(−)_ increases after surprising no reward outcomes. This learning rate was not allowed to be less than 0, such that

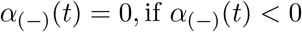

To generate choice probabilities, the *Q*-values were fed into the same softmax decision function as the static-learning model.

We also examined two other meta-learning models from the *Q*-learning family of RL models. The first is an updated form of the opponency model (Daw et al., 2002) referred to as the global reward state model (Wittmann et al., 2020). In this model, a global reward history variable influences learning from rewards and no rewards asymmetrically, as those outcomes carry different amounts of information depending on the richness of the environment. In this model, the value of a chosen action, for example *Q*_*l*_, is updated according to

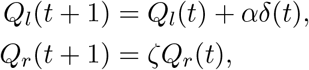

while the unchosen action value, *Q*_*r*_ is forgotten with rate *ζ*. The prediction error, *δ*, is calculated by

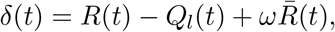

where *R* is the outcome, 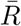 is a global reward history term and *ω* is a weighting parameter that can be positive or negative. 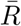 is updated on each trial:

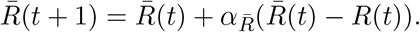

Here, 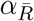 is the learning rate for the global reward term. The learned action values are converted into choice probabilities using the same softmax decision function described above.

The second model we tested is an adapted Pearce-Hall model (Pearce and Hall, 1980) in which the learning rate is a function of RPE magnitude. If the left action is chosen, *Q*_*l*_ is updated by the learning rule

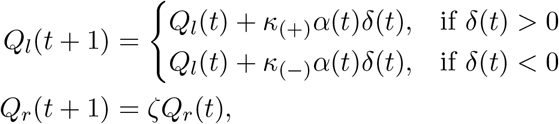

where *κ*_(+)_ and *κ*_(−)_ are the salience parameters for rewards and no rewards, respectively. Having separate salience parameters is a modification of the original model that we made to improve fit and mirror the asymmetry in our own meta-learning model and the global reward state model. The learning rate *α* is updated a function of RPE:

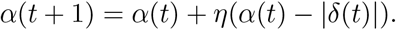

Here, *η* controls the rate at which the learning rate is updated. In this way, the model enhances learning rates when the recent average of RPE magnitudes is large. This approach contrasts with our meta-learning model which diminishes the learning rate as a result of large recent RPE magnitudes if they are consistent.

### Data analysis: firing rate model

We developed a version of our meta-learning model to fit inter-trial firing rates to see if neural activity and choice behavior reported similar dynamics of expected uncertainty. The learning components of the models were identical, but the firing rate model fit z-scored firing rates as a function of expected uncertainty:

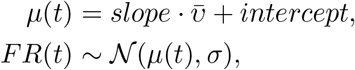

where *slope* and *intercept* scale expected uncertainty into the mean predicted firing rate, *μ*. Real firing rates, *FR*, are modeled as a draw from a Gaussian distribution with mean *μ* and some fixed amount of noise, *σ*.

### Data analysis: Model fitting

We fit and assessed models using MATLAB (Mathworks) and the probabilistic programming language, Stan (https://mc-stan.org/) with the MATLAB interface, MatlabStan (https://mc-stan.org/users/interfaces/matlab-stan). Stan was used to construct hierarchical models with mouse-level hyper-parameters to govern session-level parameters. For each session, each parameter in the model (for example, *α*_*ϵ*_ for the meta-learning model) was modeled as a draw from a mouse-level distribution with mean *μ* and variance *σ*. Models were fit using noninformative (uniform distribution) priors for session-level parameters ([0, 1] for all parameters except *β* which was [0, 10]) and weakly informative 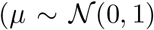, *σ* ~ half-Cauchy(0, 3)) priors for mouse-level hyperparameters. For some mice with fewer sessions, more informative mouse-level hyperparmeters were used to achieve model convergence under the assumption that individual mice behave similarly across days. This hierarchical construction mitigated the typical variability of point estimates for session-level parameters that results from other methods of estimation. Stan uses full Bayesian statistical inference to generate posterior distributions of parameter estimates using Hamiltonian Markov chain Monte Carlo sampling (Carpenter et al., 2017). The parameters for updating expected uncertainty, *α*_*v*_, and for updating the negative RPE learning rate, *ψ*, were ordered such that *ψ > α*_*v*_. The ordering operated under the assumption that learning rate should be integrated more quickly to detect change. The ordering also helped models to converge more quickly.

### Data analysis: extracting model parameters and variables, behavior simulation

For extracting model variables (like expected uncertainty), we took at least 4,000 draws from the Hamiltonian Markov Chain Monte Carlo samples of session-level parameters, ran the model agent through the task with the actual choices and outcomes, and averaged each model variable across runs. For comparisons of individual parameters across behavioral and neural models, we obtained maximum *a posteriori* parameter values by approximating the mode of the distribution: binning the values in 50 bins and taking the median value of the most populated bin. For simulations of behavior, we took at least 4,000 draws from the Hamiltonian Markov Chain Monte Carlo samples of mouse-level parameters and simulated behavior and outcomes in a number of random sessions per sample. For the transition analysis, that number was proportional to the number of rare transitions that each animal contributed to the actual data. For other analyses that number was fixed.

### Data analysis: linear regression models of neural activity

For comparisons of firing rates to the behavioral-model-generated uncertainty terms we regressed z-scored firing rates on z-scored uncertainty using the MATLAB function “fitlm”. For some neurons and sessions, firing rates and model variables demonstrated monotonic changes across the session. To control for the effect of these dynamics in comparisons of inter-trial interval firing rates to model variables, we regressed out the monotonic effects for each term separately, then regressed the firing rate residuals on the expected uncertainty residuals. Here, we found similar rates of correlation across the population of neurons. We also looked for relationships between the neural activity and other model variables that evolved as a function of action and outcome history. For the analysis of the dynamic foraging task data, we added total value (*Q*_*r*_ + *Q*_*l*_), relative value (*Q*_*r*_ − *Q*_*l*_), value confidence (|*Q*_*r*_ − *Q*_*l*_|), RPE, and reward history as regressors in the same model (Figure A3e). Value confidence captures how much better the better option is on each trial. Reward history is an arbitrarily smoothed history of all rewards, generated by convolving rewards with a recency-weighted kernel. The kernel was derived from an exponential fit to the coefficients from the regression of choices on outcomes. For the dynamic Pavlovian task data, we added RPE and reward history as regressors (Figure A4e).

### Data analysis: linear mixed effect models

To analyze the changes in transition behavior we constructed a linear mixed effects model that predicted choice averages after transition points as a function of trial since transition, transition type, and the interaction between the two. The model is described by the following Wilkinson notation:

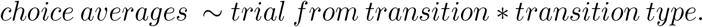

For assessing the affect of chemogenetic manipulation, we added drug condition (vehicle or agonist 21) as a fixed effect as well as the interaction between transition type and drug condition:

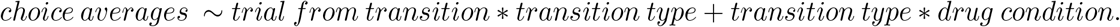

In the case of simulated data, these fixed effects were grouped by mouse, treated as a random effect that affects both slope and intercept, given by:

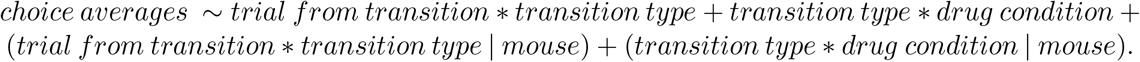

In all models, we z-scored all choice probabilties to center the data.

### Data analysis: autocorrelation controls

To control for potential statistical confounds in correlating two variables with similar autocorrelation functions — in particular, firing rates of serotonin neurons and dynamics of expected uncertainty — we simulated each variable and compared it to the real data. We simulated 1,000 expected uncertainty variables by using maximum *a posteriori* parameter estimates to simulate a random sequence of choices and outcomes of the same length as the real session. For each simulation we extracted model variables using the sampling and averaging method described above. Linear regressions of real firing rates on each simulated variable were performed. If the *t*-statistics from the regression of real firing rate on real model variable fell beyond the 95% boundary of the distribution of *t*-statistics from the comparisons with simulated variables, then the relationship was deemed significant. We view this control analysis as an estimate of a lower bound on the true rate of correlated variables; for example, in a recent paper, only approximately one-third of true correlations were recoverable with this simulation (Elber-Dorozko and Loewenstein, 2018).

Conversely, we simulated neural data with autocorrelation functions matched to those of the actual neuron. For each neuron, we computed the autocorrelation function for lags of 10 trials and calculated the sum. The autocorrelation function sum was mapped onto the scale of a half-Gaussian smoothing kernel (width of 10 trials) using a log transformation. Neurons were then simulated as a random walk such that the firing rate at a given trial was the sum of the previous 10 trials weighted by the smoothing kernel plus some normally distributed noise 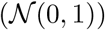. We found that the autocorrelation functions and the distributions of simulated firing rates were similar to those of the real neurons. For each real neuron, we performed 1,000 simulations and compared them to the real expected uncertainty in the same way as described above.

### Histology

After experiments were completed, mice were euthanized with an overdose of isoflurane, exsanguinated with saline, and perfused with 4% paraformaldehyde. The brains were cut in 100-*μ*m-thick coronal sections and mounted on glass slides. We validated expression of rAAV5-EF1a-DIO-hChR2(H134R)-EYFP, pAAV-hSyn-DIO-hM4D(Gi)-mCherry, or pAAV-hSyn-DIO-mCherry with epifluorescence images of dorsal raphe (Zeiss Axio Zoom.V16). In electrophysiological experiments, we confirmed targeting of the optic-fiber-tetrode bundle to the dorsal raphe by location of the electrolytic lesion.

---

**Figure A1.**
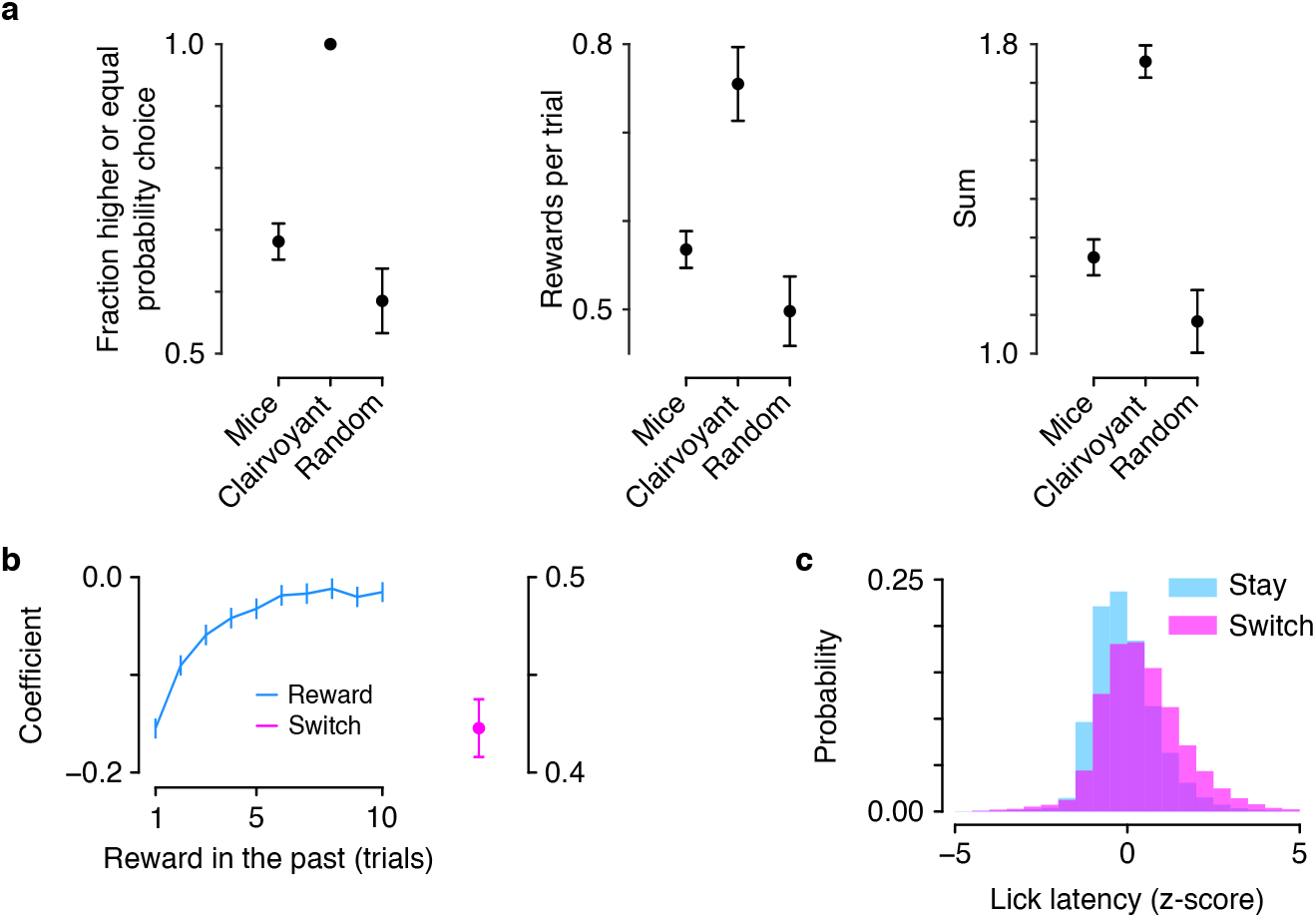
(a) Fraction of higher-probability choices, rewards per trial, and the sum of these quantities for mice, a “clairvoyant” model that knew reward probabilities, and random choices (paired *t*-test between mice and random: higher-probability choice, *t* = 11.11, *p <* 10^−18^; rewards per trial, *t* = 10.87, *p <* 10^−17^; sum, *t* = 12.30, *p <* 10^−20^). (b) Linear regression coefficients of response time on reward history. Coefficient for switch trials was included in the regression. (c) Lick latency was faster on trials in which mice repeated the same choice (“stay”) compared to when they made a different choice (“switch”; paired *t*-test, *t*_47_ = −12.85, *p <* 10^−16^).

**Figure A2.**
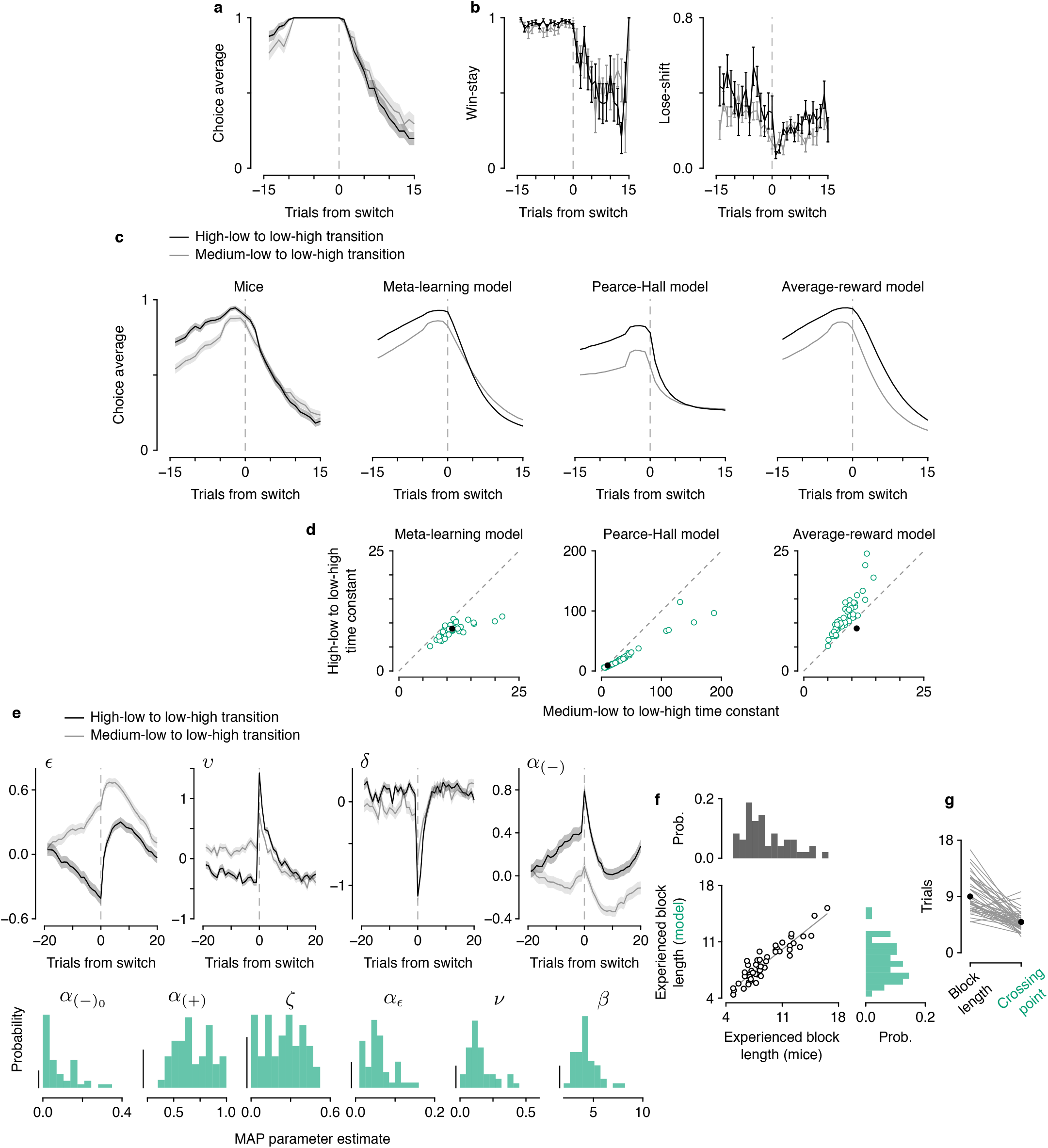
Meta-learning model: data and model comparisons. (a) Mice showed variable learning rates even when choice history leading up to transitions was identical. (b) Probability of repeating a rewarded choice (“win-stay”) and switching following an unrewarded choice (“lose-shift”) around transitions. (c) Choice averages (relative to initially-higher spout) for mice and our meta-learning model (both reproduced from Figure 2b), compared to two other models with variable learning rates: Pearce-Hall and an average-reward model. (d) The meta-learning model (green) captures transition behavior (black; panel reproduced from Figure 2e), whereas the other two models with variable learning rates do not. (e) Top: trial-by-trial dynamics of expected uncertainty (*ϵ*), unexpected uncertainty (*v*), reward prediction error (*δ*), and negative learning rate (*α*_(−)_) around transitions in reward probabilities (cf. Figure 2b). Mean *±* S.E.M. z-scored values are plotted for each variable. Bottom: maximum *a posteriori* (MAP) parameter estimates. Scale bars: 0.1. (f) Experienced block lengths were similar between mice and models. (g) The number of trials it took the model to discriminate the expected uncertainty in high compared to medium blocks (“crossing point”) was less than the experienced block length. Ignores 8 mice that did not distinguish within 30 trials or distinguished before block beginning.

**Figure A3.**
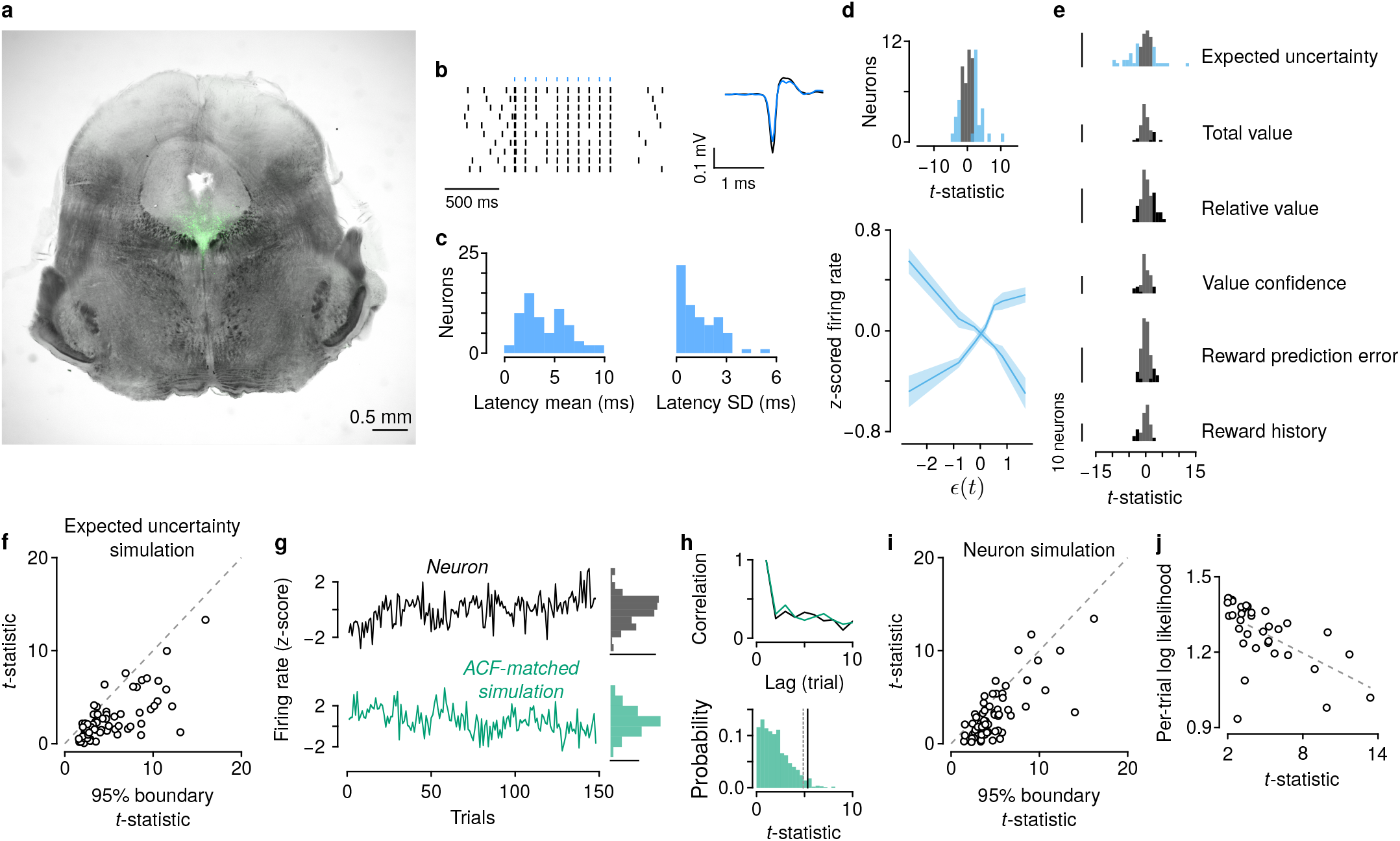
Serotonin neuron firing rates correlate with expected uncertainty. (a) Representative histological section of the midbrain from electrophysiological experiments, showing ChR2-EYFP expression (green) in dorsal raphe serotonin neurons. (b) Example of identified serotonin neuron firing in response to most light stimuli activating ChR2 with short latency and similar extracellular action potential waveform. (c) Mean and SD of firing latency of identified serotonin neurons. (d) Regression results as in Figure 3d,e, removing slow, session-long trends. (e) Distributions of *t*-statistics of regressors in a multivariate generalized linear model of inter-trial interval firing rate. (f) *t*-statistics from neurons compared with true and simulated expected uncertainty. (g) Example simulated neuron with an autocorrelation function (ACF) matched to the real neuron. Probability density scale bars: 0.2. (h) Top: ACF matching between real neuron and simulations. Bottom: distribution of *t*-statistic from the real neuron (black line) and simulations (green). Dashed gray line shows 95% boundary from the distribution of simulations. (i) *t*-statistics from real and simulated neurons compared with expected uncertainty. (j) Success of firing rate model fit correlates with *t*-statistic comparing firing rate to behavior-model-derived expected uncertainty (*R*^2^ = 0.34, Spearman’s *ρ* = −0.65, *p <* 10^4^).

**Figure A4.**
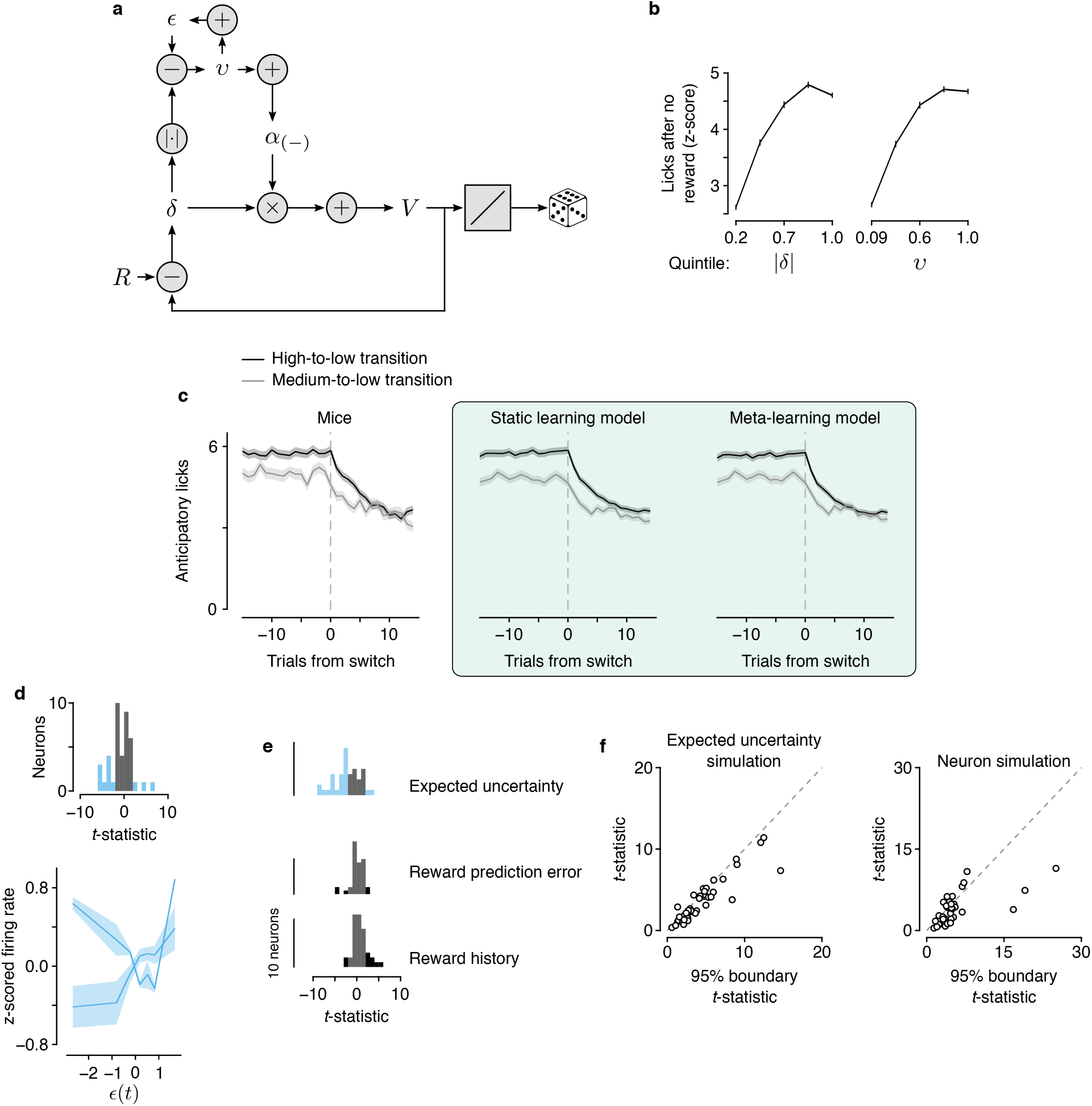
Serotonin neuron firing rates correlate with expected uncertainty in a dynamic Pavlovian task. (a) Schematic of meta-learning model applied to behavior in the dynamic Pavlovian task. The value (*V*) of the stimulus is updated analogously to the way action values (*Q*_*l*_ and *Q*_*r*_) are updated in the dynamic foraging task. *V* is mapped to licks through a linear scaling and sampling from a Poisson distribution. (b) Lick rate after no reward scales with unsigned RPE (|*δ*|, regression coefficient = 0.68, *R*^2^ = 0.11) and unexpected uncertainty (*ϵ*, regression coefficient = 0.51, *R*^2^ = 0.083). (c) Transition behavior in the dynamic Pavlovian task when probabilities changed from high to low or medium to low. Left: mice. Middle: static learning model simulated behavior. Right: meta-learning model simulated behavior. (d) Regression results as in Figure 4e,f, removing monotonic, session-long trends. (e) Distributions of *t*-statistics of regressors in a multivariate generalized linear model of inter-trial interval firing rate. (f) Left: *t*-statistics from neurons compared with true and simulated expected uncertainty. Right: *t*-statistics from real and simulated neurons compared with expected uncertainty.

**Figure A5.**
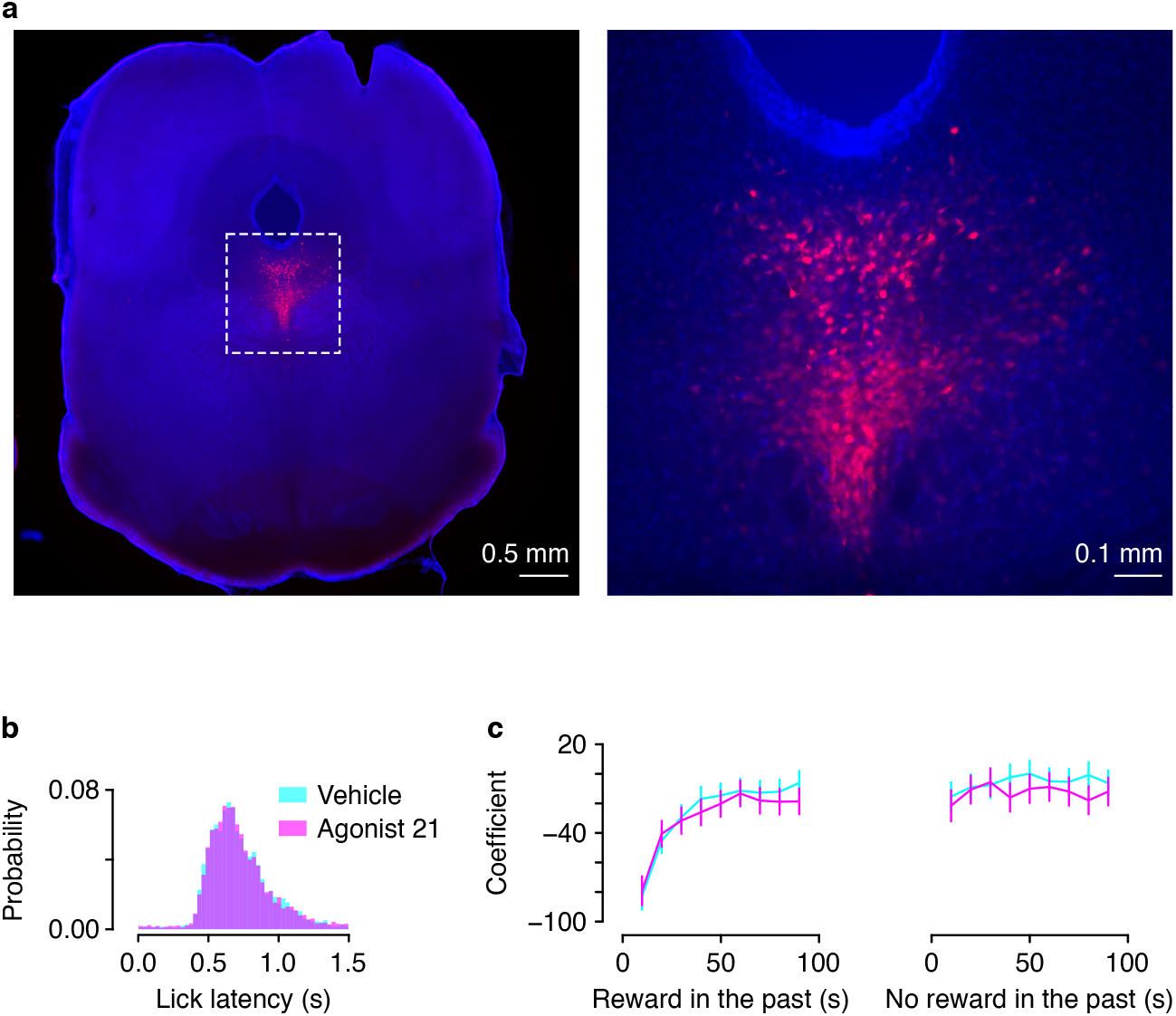
(a) Representative histological section showing DREADD expression in dorsal raphe serotonin neurons (reproduced from Figure 5a). Dashed box in the left image indicates higher-magnification image on the right. (b) No difference in lick latency comparing vehicle injections to agonist 21 injections (paired *t*-test, *t*_2_ = −0.39, *p* = 0.73). (c) Regression coefficients modeling lick latency as a function of reward (left) or no reward (right) in the past, for vehicle or agonist 21 injections.

